# Genomic variants among threatened *Acropora* corals

**DOI:** 10.1101/349910

**Authors:** S. A. Kitchen, A. Ratan, O. C. Bedoya-Reina, R. Burhans, N. D. Fogarty, W. Miller, I. B. Baums

## Abstract

Genomic sequence data for non-model organisms are increasingly available requiring the development of efficient and reproducible workflows. Here, we develop the first genomic resources and reproducible workflows for two threatened members of the reef-building coral genus *Acropora*. We generated genomic sequence data from multiple samples of the Caribbean *A. cervicornis* (staghorn coral) and *A. palmata* (elkhorn coral), and predicted millions of nucleotide variants among these two species and the Pacific *A. digitifera*. A subset of predicted nucleotide variants were verified using restriction length polymorphism assays and proved useful in distinguishing the two Caribbean Acroporids and the hybrid they form (“*A. prolifera*”). Nucleotide variants are freely available from the Galaxy server (usegalaxy.org), and can be analyzed there with computational tools and stored workflows that require only an internet browser. We describe these data and some of the analysis tools, concentrating on fixed differences between *A. cervicornis* and *A. palmata*. In particular, we found that fixed amino acid differences between these two species were enriched in proteins associated with development, cellular stress response and the host’s interactions with associated microbes, for instance in the Wnt pathway, ABC transporters and superoxide dismutase. Identified candidate genes may underlie functional differences in the way these threatened species respond to changing environments. Users can expand the presented analyses easily by adding genomic data from additional species as they become available.

**Article Summary:** We provide the first comprehensive genomic resources for two threatened Caribbean reef-building corals in the genus *Acropora*. We identified genetic differences in key pathways and genes known to be important in the animals’ response to the environmental disturbances and larval development. We further provide a list of candidate loci for large scale genotyping of these species to gather intra- and interspecies differences between *A. cervicornis* and *A. palmata* across their geographic range. All analyses and workflows are made available and can be used as a resource to not only analyze these corals but other non-model organisms.

## INTRODUCTION

Genomic data for non-model organisms are becoming available at an unprecedented rate. Analyses of these data will advance our understanding of the capacity of organisms to adapt, acclimatize or shift their ranges in response to rapid environmental change (Savolainen *et al.* 2013). While genome sequencing itself has become routine, bioinformatics treatment of the data still presents hurdles to the efficient and reproducible use of this data (Nekrutenko and Taylor 2012). Thus, genomic variant analysis workflows (e.g. Bedoya-Reina et al. (2013) are needed to eliminate some of these computational hurdles and increase reproducibility of analyses. Here, we develop such tools, apply them to threatened reef-building corals and present novel findings with respect to the molecular pathways used by these species to respond to environmental stimuli.

The *Acropora* species, *A. cervicornis* and *A. palmata* were the main reef-building corals of the Caribbean (Figure 1). These corals have greatly decreased in abundance during recent years due to infectious disease outbreaks, habitat degradation, storm damage, coral bleaching, outbreaks of predators and anthropogenic activities (Bruckner 2002). A large body of previous studies has investigated the effects of environmental stress in Caribbean Acroporid corals (Randall & Szmant 2009; DeSalvo *et al.* 2010; Baums *et al.* 2013; Libro *et al.* 2013; Polato *et al.* 2013; Parkinson *et al.* 2015). These studies highlight changes in the molecular, cellular, and physiological response of these species to an unprecedented elevation in seawater temperature. Increases in water temperature of only 2 -3 °C can reduce the fertilization rates, reduce larval survival, and deplete genotypic diversity of Caribbean Acroporids (Randall & Szmant 2009; Williams & Miller 2012; Baums *et al.* 2013).

**Figure 1.**
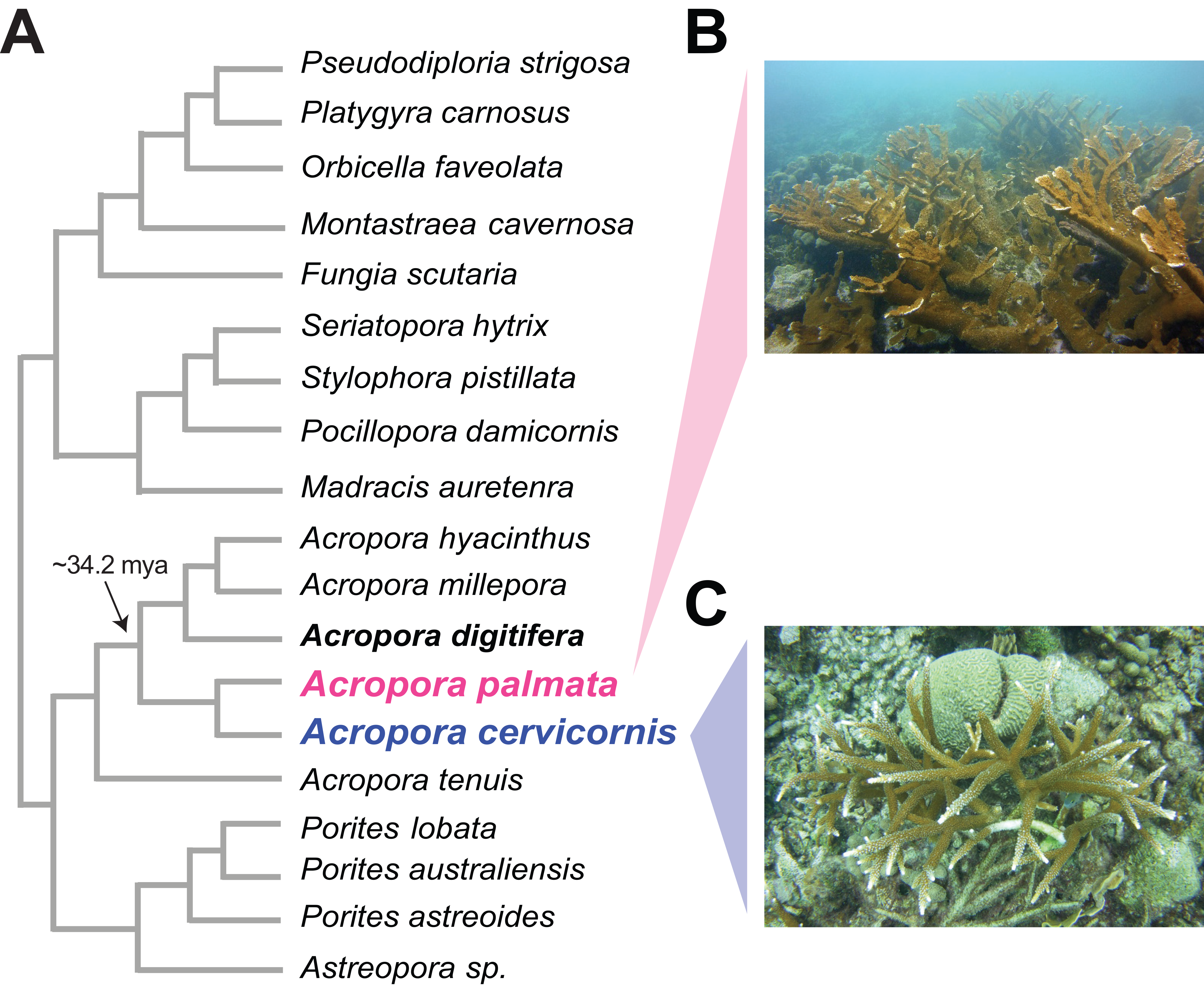
Phylogeny of corals with genomic and transcriptomic resources used in this study (A) with images of the two focal species, *Acropora palmata* (B) and *Acropora cervicornis* (C). The evolutionary relationships depicted in the coral phylogeny are redrawn based on the phylogenomic analysis by Bhattacharya et al. (2016), but branch lengths do not reflect evolutionary distance. Estimate of divergence time between the Caribbean Acroporids and *A. digitifera* was calculated by Richards et al. (2013). Photographs of *A. palmata* (B) and *A. cervicornis* (C) were taken by Iliana B. Baums (Curacao 2018).

Because of a tremendous die-off, both species are now listed as threatened on the United States Federal Endangered Species List (Anonymous 2006). Extensive conservation efforts are currently underway across the range, which will be considerably facilitated by the acquisition of genomic data. For instance, these data will help to identify management units, evolutionary significant units, hybridization dynamics, genotypic diversity cold-spots and interactions with the corals’ obligate symbionts in the genus *Symbiodinium* (Baums 2008; van Oppen et al. 2015). The project described here represents an early effort to move beyond low-resolution sequencing and microsatellite studies (Vollmer & Palumbi 2007; Baums *et al.* 2014) and employ the power of full-genome analysis (Drury et al. 2016).

Here, we present genome-wide single nucleotide variants (SNVs) between the two Caribbean Acroporids relying on the genome assembly for a closely related species, *A. digitifera* (Shinzato et al. 2011) (Figure 1). We have successfully used the same approach to analyze genomes using much more distant reference species, such as polar, brown and black bears based on the dog genome (Miller *et al.* 2012), and giraffe based on cow and dog (Agaba et al. 2016). We highlight several examples of how these SNVs enable population genomic and evolutionary analyses of two reef-building coral species. The SNV results are available on the open source, public server Galaxy (Afgan et al. 2016), along with executable histories of the computational tools and their settings. This workflow presented here for corals and by Bedoya-Reina et al. (2013) can be transferred for genomic analyses of other non-model organisms, and provide abundant information in a reproducible manner.

## MATERIALS AND METHODS

### DNA Extraction and Sequencing

For each species, five previously genotyped samples from the Baums Lab coral tissue collection were selected from each of the four sites representing their geographic range: Florida (FL), Belize (BE), Curacao (CU) and U.S. Virgin Islands (VI; Table 1) (Baums et al. 2009; Baums et al. 2005). An additional sample for each species from Florida (*A. cervicornis* CFL14120 and *A. palmata* PFL1012) was selected for deep genome sequencing because they are located at easily accessible and protected sites in the Florida Keys (*A. palmata* at Horseshoe Reef and *A. cervicornis* at the Coral Restoration Foundation nursery), and are predictable spawners that are highly fecund. High molecular weight DNA was isolated from each sample using the Qiagen DNeasy kit (Qiagen, Valencia, CA) according to the manufacturer’s protocol. DNA quality and quantity was assessed with gel electrophoresis and Qubit 2.0 fluorometry (Thermo Fisher, Waltham, MA), respectively. Sequence library construction and sequencing was completed by the Pennsylvania State University Genomics Core Facility. Paired-end short insert (550 nt) sequencing libraries of the two deeply sequenced genomes were constructed with 1.8-2 μg sample DNA and the TruSeq DNA PCR-Free kit (Illumina, San Diego, CA). The remaining 40 paired-end short insert (350 nt) sequencing libraries (Table S1) were constructed using 100 ng sample DNA and the TruSeq DNA Nano kit (Illumina, San Diego, CA). Deep- and shallow- sequence libraries were pooled separately and sequenced on the Illumina HiSeq 2500 Rapid Run (Illumina, San Diego, CA) over two lanes and four lanes, respectively.

**Table 1.**
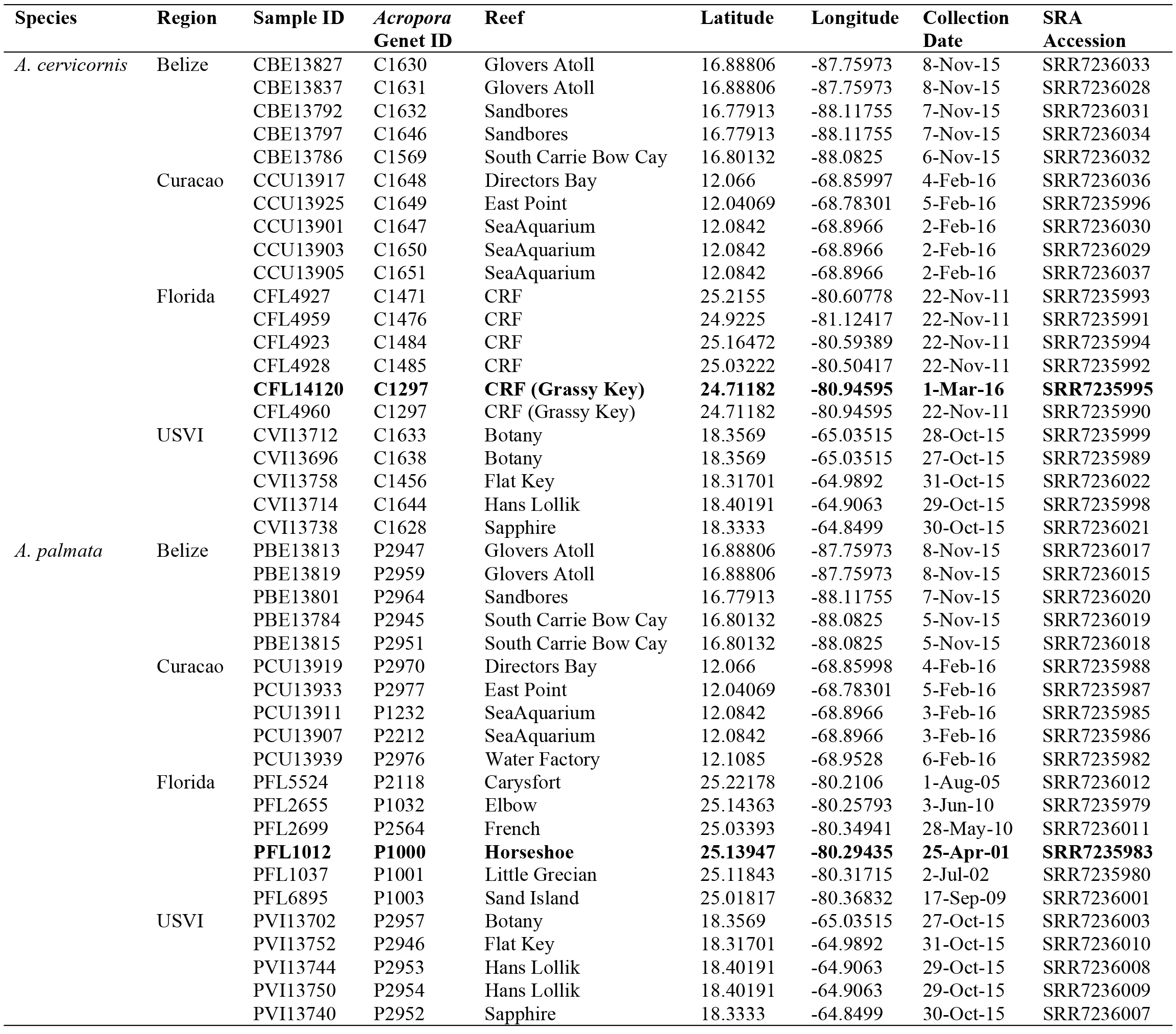
Sequenced Genomes. Species assignment was based initially on microsatellite multilocus genotyping. *Acropora* Genet ID is an identifier for each *Acropora* multilocus microsatellite genotype in the Baums Lab database. Coordinates are given in decimal degrees (WGS84). Two samples were sequenced to a greater depth (bold type).

### *A. digitifera* Assembly and Inter-species Gene Model Comparisons

We downloaded the *A. digitifera* genome assembly and GFF-formatted gene annotations from NCBI GCA_000222465.2 Adig_1.1). To conduct the pathway enrichment analysis, we obtained additional annotation from the Kyoto Encyclopedia of Genes and Genomes (KEGG) (Kanehisa et al. 2017). During gene prediction, gene annotation can be error prone and misled by assembly gaps or errors, imprecision of *de novo* gene predictors and/or errors in gene annotations in the species used for comparison, among other sources. To overcome these known issues, our approach included, at a minimum, submitting the putative amino acid sequence to the blastp server maintained by the Reef Genomics Organization (Liew et al. 2016) () and the blastp and/or psi-blast servers at NCBI (Altschul et al. 1997) (). We also used the Reef Genomics website to assess the degree of inter-species sequence conservation among 20 corals in Figure 1 (resources include transcriptomes and genomes, details provided in Bhattacharya et al. (2016), and the Genome Browser () at the University of California at Santa Cruz (Kent et al. 2002) to measure the inter-species conservation of the orthologous mammalian residue. We interpret the degree of conservation at a protein position and its immediate neighbors as suggesting the amount of selective pressure and the functional importance of the site.

### Single Nucleotide Variant and Indel Calls

We aligned the paired-end sequences for the 42 samples to the *A. digitifera* reference genome sequence using BWA version 0.7.12 (Li and Durbin 2009) with default parameters. On average, we were able to align ~89% of the reads for each individual, and ~74% of the reads aligned with a mapping quality > 0. Paired-end reads are generated by sequencing from both ends of the DNA fragments, and we found that about 70% of these reads aligned within the expected distance from its mate in those alignments (see Table S1 for details). We used SAMBLASTER version 0.1.22 (Faust and Hall 2014) to flag potential PCR duplicate reads that could otherwise affect the quality of the variant calls (Table S1). Considering data from all individuals simultaneously, we used SAMtools version 1.3.1 (Li et al. 2009) to identify the locations of putative variants with parameters ‒g to compute genotype likelihoods, -A to include all read pairs in variant calling, and ‒E to recalculate the base alignment quality score against the reference *A. digitifera* genome. Variants were called with bcftools version 1.2 (Li 2011) multiallelic caller and further filtered to keep those variants for which the total coverage in the samples was less than 1,200 reads (to limit the erroneous calling of variant positions in repetitive or duplicated regions), the average mapping quality was greater than 30, and the fraction of reads that aligned with a zero mapping quality was less than 0.05. The VCF file of nucleotide variants was converted to gd_snp format using the “Convert” tool from the “Genome Diversity” repository on Galaxy, after separating the substitution and insertion/deletion (indel) variants. The resulting sets of SNVs and indels are available on Galaxy (*a permanent link will be made available upon acceptance*). The mitochondrial variants were similarly identified using the *A. digitifera* mitochondrial reference genome (GenBank: NC_022830), and Figure S3 was drawn using Millerplot (https://github.com/aakrosh/Millerplot).

The Galaxy tool “Phylogenetic Tree” under Genome Diversity (Bedoya-Reina et al. 2013) was used to calculate the genetic distance between two individuals at a given SNV as the difference in the number of occurrences of the first allele. For instance, if the two genotypes are 2 and 1, i.e., the samples are estimated to have respectively 2 and 1 occurrences of the first allele at this location, then the distance is 1 (the absolute value of the difference of the two numbers). The Neighbor-joining tree was constructed with QuickTree (Howe et al. 2002) and visualized with draw_tree utility script in package PHAST (Hubisz et al. 2010). We used I-TASSER online server for protein structure prediction (Yang et al. 2015) to model and further help to develop hypotheses about functionality of several mutations in STE20-related kinase adapter protein alpha protein (NCBI: L0C107340566). Identification of enriched KEGG pathways was completed using the “Rank Pathways” tool, which compares the gene set with SNVs against the complete set of genes in the pathway using the statistical Fisher’s exact test.

### Genomic Regions of Differentiation

We assigned a measure of allele frequency difference to each SNV analogous to calculations of *F*_ST_ for intra-species comparisons using the “Remarkable Intervals” Galaxy tool (score shift set to 90%). *F*_ST_ values can be used to find genomic regions where the two species have allele frequencies that are remarkably different over a given window or interval, i.e., the *F*_ST_ s are unusually high. Such intervals may indicate the location of a past “selective sweep” (Akey et al. 2002) caused by a random mutation that introduces an advantageous allele, which rises to prominence in the species because of selective pressures, thereby increasing the frequency of nearby variants and changing allele frequencies from those in an initially similar species. In theory, the *F*_ST_ ranges between 0, when the allele frequencies are identical in the two species, to 1, for a fixed difference. However, in practice it works better to use an estimation formula that accounts for the limited allele sampling; we employ the “unbiased estimator” of Reich et al. (2009), because it performs best on the kinds of data used here, according to Willing *et al.* (2012). It should be noted that care must be taken when interpreting high *F*_ST_ values this way, since they can also be caused by genetic drift, demographic effects, or admixture (Holsinger and Weir 2009). We compared these intervals to the genome-wide *F*_ST_ estimate calculated using the Galaxy tool “Overall FST”.

### PCR-Ready SNV Markers and RFLP Validation

PCR-ready SNVs were identified based on the following criteria: 1) the SNV-caller considered them to be high-quality (Phred-scaled quality score ≥ 900), 2) all 21 *A. cervicornis* samples looked homozygous for one allele while all 21 *A. palmata* samples looked homozygous for the other allele and 3) there were no observed SNVs, indels, low-complexity DNA or unassembled regions within 50 bp on either side of the SNV.

From the PCR-ready SNVs, we developed a PCR-restriction fragment length polymorphism (RFLP) assay to validate a subset of fixed SNVs with additional Caribbean Acroporids samples, including the hybrid of the two species, *Acropora prolifera* (Table S2). We screened 197 fixed SNVs with 50bp flanking sequence (101bp total) using the webserver SNP-RFLPing2 (Chang et al. 2006; Chang et al. 2010) to find a set of loci that would cut with common restriction enzymes (*Hae*III, *Dpn*II, *Hinf*I, *Eco*RV, and *Hpy*CH4IV all from New England Biolabs, Ipswich, MA). Eight loci were selected, of which half cut *A. palmata*-like SNVs while the other half cut *A. cervicornis*-like SNVs (Table S3). For each diagnostic locus, additional flanking sequence was extracted from the scaffold until another restriction enzyme recognition site was encountered for that specific locus-restriction enzyme combination. Primers were designed for the extended flanking sequence using Primer3web version 4.1.0 (Untergasser et al. 2012).

A reference set of parental (*n*= 10 *A. palmata* and *n*= 9 *A. cervicornis*) and hybrid (*n* = 27 colonies) samples from across the geographic range were tested with a previously developed microsatellite assay based on five markers (Baums et al. 2005) and the RFLP assay (Table S2). Atest set of hybrids (*n*=20 colonies) that did not have previous genetic information was also included to compare taxon assignment between the two marker sets. Hybrids were initially identified in the field based on intermediate morphological features following Cairns (1982); Van Oppen et al. (2000); Vollmer and Palumbi (2002).

For all samples, DNA was extracted using the DNeasy kit (Qiagen, Valencia, CA). PCR reactions consisted of 1X NH_4_ Buffer (Bioline, Boston, MA), 3 mM MgCl_2_ (Bioline, Boston, MA), 1 mM dNTP (Bioline, Boston, MA), 250 nmol forward and reverse primers (IDT, Coralville, Iowa), 1 unit of Biolase DNA polymerase (Bioline, Boston, MA) and 1 μl of DNA template for a total volume of 10μl. The profile for the PCR run was as follows: 94 °C for 4 min for initial denaturing, followed by 35 cycles of 94 °C for 20s, 55 °C for 20s, and 72 °C for 30s, and a final extension at 72 °C for 30min. For each locus, 5 μl of PCR product was combined with 1X restriction enzyme buffer (New England Biolabs, Ipswich, MA) and 0.2 μl restriction enzyme (New England Biolabs, Ipswich, MA) for a total reaction volume of 10 μl and incubated according to the manufacturer’s recommendation. PCR and digest fragment products were resolved by 2% TAE agarose gel electrophoresis at 100 V for 35 min, except for locus NW_015441368.1: 282878 that was run on 3.5% TAE agarose gel at 75 V for 45 min to resolve the smaller fragments. Banding patterns were scored for each locus as homozygous for either parent species (1 or 2 bands) or heterozygous (3 bands).

Reference samples were first assigned to taxonomic groups (*A. palmata*, *A. cervicornis*, F1 or later generation hybrid) based on allele frequencies at five microsatellite loci (Baums et al. 2005) by NEWHYBRIDS (Anderson and Thompson 2002).A discriminant factorial correspondence analysis (DFCA) was performed on the microsatellite and SNV marker data separately to predict sample membership to the taxonomic groups: *A. palmata*, *A. cervicornis*, F1 11hybrid or later generation hybrid. The FCA performed in GENETIX version 4.05 (Belkhir et al. 2004) clustered the individuals in multi-dimensional space based on their alleles for each marker type. The factorial axes reveal the variability in the data set with the first factor being the combination of alleles that accounts for the largest amount of variability. The FCA scores for all axes were used ina two-step discriminant analysis using the R statistical software (RCoreTeam 2017) to calculate the group centroid, or mean discriminant score for a given group, and individual probability of membership to a given group using leave-one-out cross-validation First, the parameter estimates for the discriminant function of each group were trained by the FCA scores from the reference samples. Second, those functions were used to assign all samples, including the test set of hybrids, based on their FCA scores to a taxon group.

### Data Availability

The executable histories for the SNV and protein analyses and their respective data sets are available on Galaxy (link provided in final publication). Table 2 lists the data sets available on Galaxy. Specifically, the data sets “coral snps” and “intra-codon variants” are tables of variants with positions in reference to the *A. digitifera* genome. The data set “PCR-Ready SNVs” are 101 bp sequences extracted from the *A. digitifera* genome, with 50 bp flanking sequence surrounding the fixed SNV. Raw sequence data are deposited in the NCBI Sequence Read Archive (accessions SRR7235977-SRR7236038).

**Table 2.** Data sets available on Galaxy.

| Name | Contents | # of Lines |
| --- | --- | --- |
| SNVs | *A. digitifera* scaffold positions with two observed nucleotides among the three *Acropora* genomes | 8,368,985 |
| indels | positions and contents of observed short ($\leq 20$ bp) insertion/deletions | 940,345 |
| SAPs | protein sequence positions of non-synonymous and synonymous substitutions | 561,015 |
| mitochondrial SNVs | *A. digitifera* mitochondrial genome positions with two observed nucleotides | 172 |
| mitochondrial indels | position of an insertion/deletion | 1 |
| exons | scaffold positions of annotated exon endpoints | 222,156 |
| PCR-ready SNVs | SNVs where no other SNV, indel, or low-complexity sequence is within 50 bp | 894 |

## RESULTS

### Variants between Three Acroporid Species

For each species, we performed deep-coverage sequencing (roughly 150-fold coverage) of one sample and shallow sequencing (roughly 5-fold to 10-fold) of 20 samples, five each from four geographic locations (Florida, the Virgin Islands, Belize, and Curacao) (Figure 2A). For details, see Table 1. The sequence coverage distribution for the Acroporid samples was comparable between species (*A. cervicornis*: Figure S1 and *A. palmata*: “coral SNPs” history at *link to be provided in final publication*).

**Figure 2.**
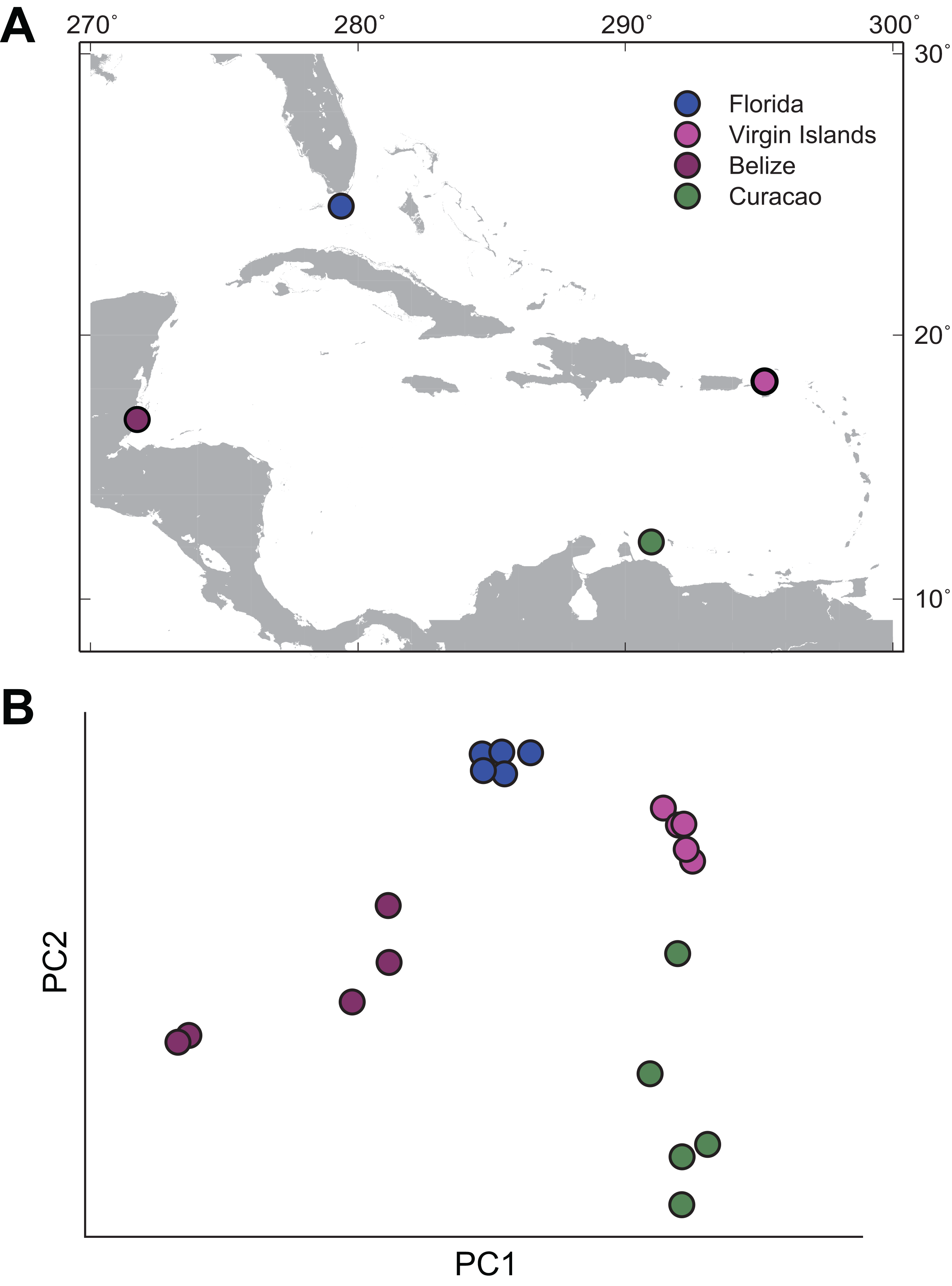
Geographic origin of *Acropora* samples (A) and Principal Components Analysis of *A. cervicornis* samples, five from each of four locations (B). As noted in analyses of other datasets (e.g., Novembre et al. (2008)) the geographic map is similar to the PCA.

Rather than relying on *de novo* assembly and gene annotation of our data, we based the analysis reported below on an assembly and annotation of the highly similar reference genome of *A. digitifera* (NCBI: GCA_000222465.2 Adig_1.1) (Shinzato et al. 2011). This strategy increases reproducibility and leverages the work of large and experienced bioinformatics groups. Important advantages of using this third species is that we can transfer its gene annotation as well as “polarize” variants, as follows. The two sequenced species in this study diverged in the Eocene about 34.2 mya from the most recent common ancestor they share with the reference species *A. digitifera* (Figure 1) (van Oppen *et al.* 2001; Richards *et al.* 2013). Thus, with a difference observed among the *A. cervicornis* and *A. palmata* samples, the allele agreeing with *A. digitifera* can be interpreted as ancestral, and the variant allele as derived.

We identified both substitution and indel variants by aligning our paired-end sequencing reads to the *A. digitifera* assembly and noting nucleotide differences with *A. cervicornis* and *A. palmata* (Table 2). Specifically, each reported substitution variant is a position in an *A. digitifera* assembly scaffold where at least one of our sequenced samples has a nucleotide that is different from the *A. digitifera* reference nucleotide, after all the thresholds on read-depth and mapping quality as discussed in the Methods were applied. We call each of these an SNV (singlenucleotide variant) because “SNP” (single-nucleotide polymorphism) is commonly used to describe an intra-species polymorphism. These data permit comparisons among the three *Acropora* species, although this paper focuses on *A. cervicornis* and *A. palmata*, and ignores unanimous differences of the new sequences from the reference.

### Fixed differences of SNVs and Indels between *A. cervicornis* and *A. palmata*

Single nucleotide variants and indels can be used to explore either intra- or inter-species variation, using similar techniques in both cases. Of the 8,368,985 SNVs, 4,998,005 are identically fixed in *A. cervicornis* and *A. palmata*, leaving 3,370,980 variable within our two sequenced species, only 1,692,739 of which were considered high-quality (Phred-scaled quality ≥ 900, Table 2). The results reported below use this set of substitution variants. A phylogenetic tree based on the genetic distance between those SNVs clearly separates the two species, and distinguishes the samples from each species according to where they were collected in most cases (Figure S2). The same is true of a Principal Component Analysis (Figure 2). From all the SNVs, both synonymous and non-synonymous amino acid substitutions were identified from the coding sequences (Table 2). Out of the 561,015 putative protein-coding SNVs, we retained the 120,206 deemed “high quality” and variable in the two newly sequenced species. To complete our analysis, we identified 172 mitochondrial SNVs, which are highly concentrated in the gene-free “control region” (Figure S3). This region also contains the only identified indel between *A. digitifera* and the two Caribbean Acroporids (Figure S3).

The examples in most of the following sections investigate only inter-species differences, and in particular focus on fixed SNVs, i.e., locations where the 21 sequenced *A. cervicornis* samples share the same nucleotide and the 21 *A. palmata* samples share a different nucleotide. Variants were filtered so that the genotype of each shallow genome within a species would match its deeply sequenced genome. This approach identified 65,533 fixed nucleotide SNV differences and 3,256 fixed amino acid differences, spread across 1,386 genes (Table 2, see Galaxy histories “coral SNPs” and “coral proteins”). These SNVs are potentially useful for investigating the genetic causes of phenotypic differences between the two *Acropora* species. In the following, by “fixed” difference we always mean fixed between *A. cervicornis* and *A. palmata*. It should be also be noted that such variants may be simply the result of demographic process rather than the result of adaptation to different niches.

Identified indels can also be analyzed to understand genomic difference between the studied species. Filtered in a manner analogous to the SNVs (requiring “high quality” and variability in *A. cervicornis* plus *A. palmata*), the original set of 940,345 genome-wide indels (Table 2) was reduced to 149,036. Of those, 2,031 were identified as fixed between *A. cervicornis* and *A. palmata*. They provide an additional set of hints for tracking down the genetic underpinnings of inter-species phenotypic differences, since indels are often more disruptive than substitutions.

### Examples of Substitutions with Potential Protein Modifications

We scanned the list of proteins with a fixed amino acid difference (or several fixed differences) to examine more closely. One potentially interesting fixed amino acid substitution is found in superoxide dismutase (SOD), whose activity is essential for almost any organism, and particularly for corals, like *Acropora*, that harbor symbionts of the genus *Symbiodinium*. This fixed difference was identified in comparison to *A. digitifera* (NCBI: LOC107335510 or Reef Genomic: Acropora_digitifera_12779), which strongly matched (E-value 3e-85) the human manganese SOD mitochondrial protein (GenBank: NP_001309746.1; Figure S4). We observed a glutamate (E) to glutamine (Q) substitution in *A. cervicornis*, corresponding to position 2 of the *A. digitifera* orthologue (Figure S4). According to the surveyed coral sequences, the Q is fixed in a number of other corals, except for an E shared by *A. digitifera*, *A. palmata*, *A. hyacinthus*, *A. millepora* and *A. tenuis* suggesting a lineage-specific mutation (Figure S4).

Another gene, NF-kappa-B inhibitor-interacting Ras-like protein 2 (NKIRAS2; NCBI: LOC107355568 and Reef Genomics: Acropora_digitifera_6635) has two putative fixed amino acid difference in the Caribbean Acroporids (Figure S5). One, an E to aspartic acid (D) substitution, occurs in the middle of a “motif’ LGT**E**RGV⟶LGT**D**RGV that is fairly well conserved between *A. palmata* and other members of the complex corals including *Porites* spp. and *Astreopora* sp. as well as robust corals except the Pocilloporidae family (*S. pistillata* and *Seriatopora spp*.), but not with *A. cervicornis* or other Acroporids (Figure S5). Thus, this appears to be a recurrent substitution in corals. The second putative fixed amino acid difference in this gene is unique to *A. cervicornis* from the corals we surveyed. The transition is from a polar but uncharged asparagine (N) to a positively charged lysine (K) in the short motif SVDGS**N**G⟶SVDGS**K**G (Figure S5). This substitution might have consequences on the tertiary structure and function of this gene in *A. cervicornis* compared to the other Acroporids.

### Fixed Indels in Protein-Coding Regions

We also looked for fixed indels in protein-coding regions among corals compared to respective mammalian orthologues. Of the 2,031 fixed indels identified, most were not found in coding sequence with only 18 genes having a fixed indel. For closer inspection, we picked a fixed indel in STE20-related kinase adapter protein alpha (STRADα; NCBI: LOC107340566, Reef Genomics: Acropora_digitifera_13579) because it has a deletion of four-amino acids, along with two amino acid substitutions in *A. palmata*, both of which are fixed differences between the Caribbean Acroporids. It aligns well with human STRADα, isoform 4 protein NP_001003788.1 (E-value 2e-77). A blastp search of coral resources indicates that the deletion is unique to *A. palmata* (Figure S6B), although *Madracis auretenra* also has a four amino acid deletion, but shifted by three positions. This deletion in *A. palmata* is confirmed by the lack of reads mapping to the 12bp nucleotide region (Figure S7).

To determine the degree of protein modification from these differences, we positioned them on a predicted protein structure of *A. cervicornis* using I-TASSER server (Yang et al. 2015). Figure 3 illustrates the predicted configuration of the protein using as structural reference the inactive STRADα protein annotated by Zeqiraj et al. (2009). The indel occurring between *A. palmata* and *A. cervicornis* is at positions 322 to 325, and the substitutions in positions 62 and 355. In order to induce the activation of STRADα, ATP binds and induces a conformational change. In its active stage, STRADα interacts with MO25α by means of the alpha-helixes B, C and E, the beta-laminae 4 and 5, and the activation loop to further regulate liver kinase B1 (LKB1) (Zeqiraj et al. 2009). Despite the fact that neither the substitutions nor the indel are placed in the structural elements described to interact with ATP or MO25α, it is difficult to disregard their functional role with them or with LKB1.

**Figure 3.**
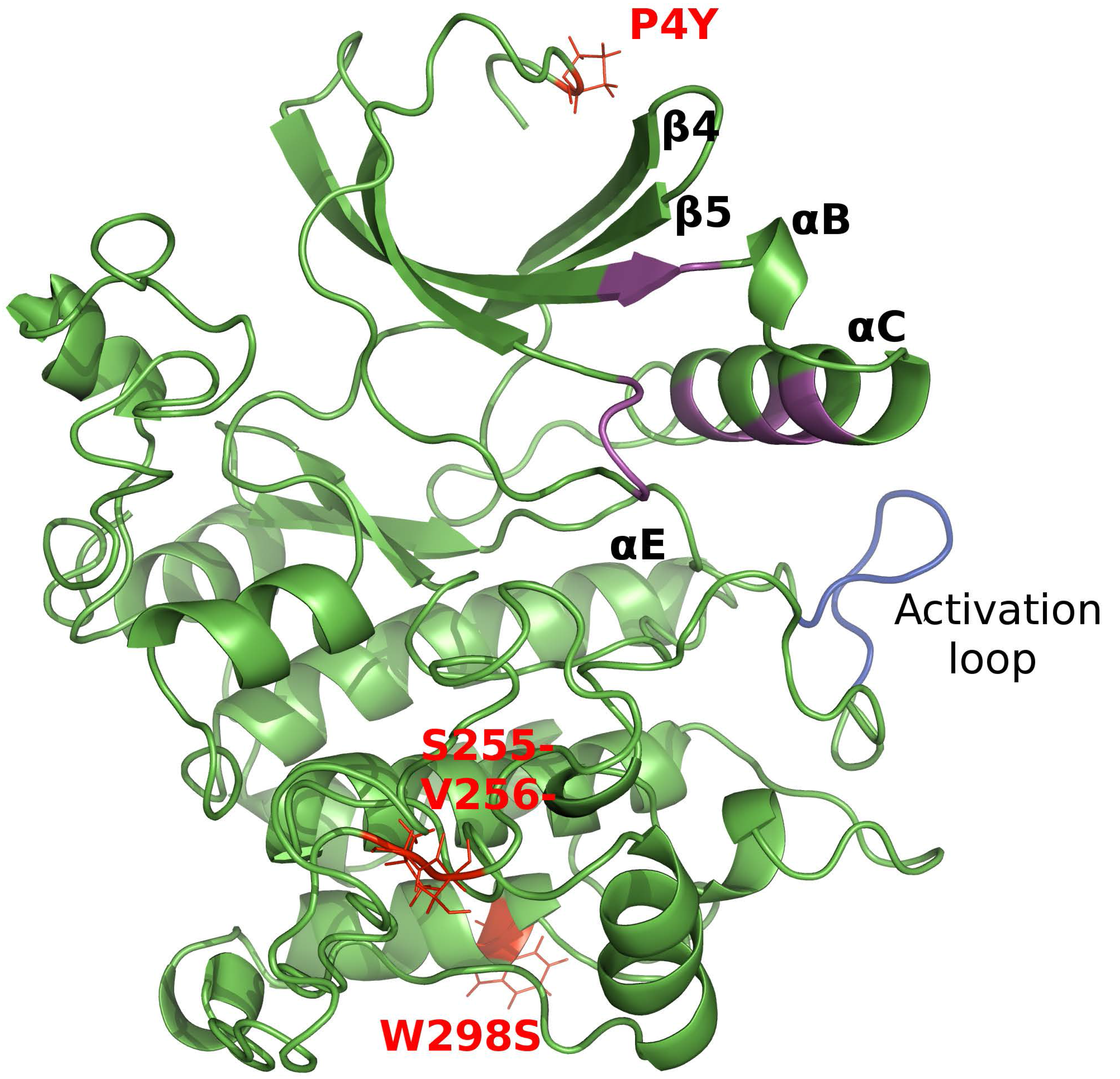
Predicted structure for STRADα in *A. cervicornis*. In its inactive conformation, ATP binds the protein to activate it (in the space delimited by the purple residues). After the protein is active, STRADα interacts with MO25 to regulate LKB1. This interaction occurs by means of the alpha-helices B, C and E, the beta-laminae 4 and 5, and the activation loop (blue). *A. palmata* differs from *A. cervicornis* in two amino acids (N62Y and P355S) as well as in four insertions (R322, D323, G324 and G325).

### KEGG Pathways Enriched for Fixed SNVs

An alternative to looking at individual amino acid substitutions is to search for protein groupings that are enriched for substitutions. This is frequently done with Gene Ontology terms (Consortium 2015) and/or classifications according to the KEGG (Kanehisa et al. 2017). We took advantage of the *A. digitifera* KEGG pathway annotations and looked for KEGG classes enriched for fixed amino acid variants. Five out of 119 pathways were found to be enriched in non-synonymous substitutions between *A. palmata* and *A. cervicornis* (two-tailed Fisher’s exact test, *p* < 0.05), and included two pathways where up to 12 genes presented these differences (i.e. ABC transporters and Wnt signaling pathway, Table 3). In Figure 4, the Wnt signaling pathway and the 12 genes with a fixed difference out of 101genes (approximately 12%) in this pathway are displayed. Note that multiple genes in Table 3 can be mapped to the same module, and several modules might appear more than once in Figure 4. In particular, these 12 genes added 27 non-synonymous fixed differences between *A. palmata* and *A. cervicornis*, and were grouped into seven different modules within the pathway (i.e. Axin, beta-catecin, Frizzled, Notum, SMAD4, SIP, and Wnt). Of these modules, Wnt grouped the largest number of genes (*n*=5), followed by Frizzled (*n*=2), and all the other modules with just one gene. The Wnt module included three WNT4 paralogue genes and nine non-synonymous mutations. Notably, the Axin module included only one gene orthologue to AXIN1 (i.e. L0C107345943) but six non-synonymous mutations. Similarly, the module Notum only includes one gene orthologue to NOTUM but this gene has five non-synonymous fixed mutations between *A. palmata* and *A. cervicornis*.

**Table 3.** Statistically significant KEGG pathways enriched for genes having a fixed amino acid difference between *A. cervicornis* and *A. palmata*. The third column gives the number of genes in the pathway with one or more fixed difference(s), and the third reports what fraction they represent of all genes in the pathway. For instance, 67 of the genes are annotated as belonging to the ABC transporter pathway, and 12/67 = 0.18. Statistical significance determined using a two-tailed Fisher’s exact test.

| Pathway | p-value | # Genes | Fraction |
| --- | --- | --- | --- |
| adf02010=ABC transporters | 0.0015 | 12 | 0.18 |
| adf00790=Folate biosynthesis | 0.019 | 5 | 0.21 |
| adf03420=Nucleotide excision repair | 0.031 | 7 | 0.15 |
| adf04933=AGE-RAGE signaling pathway in diabetic complications | 0.023 | 9 | 0.14 |
| adf04310=Wnt signaling pathway | 0.037 | 12 | 0.12 |

**Figure 4.**
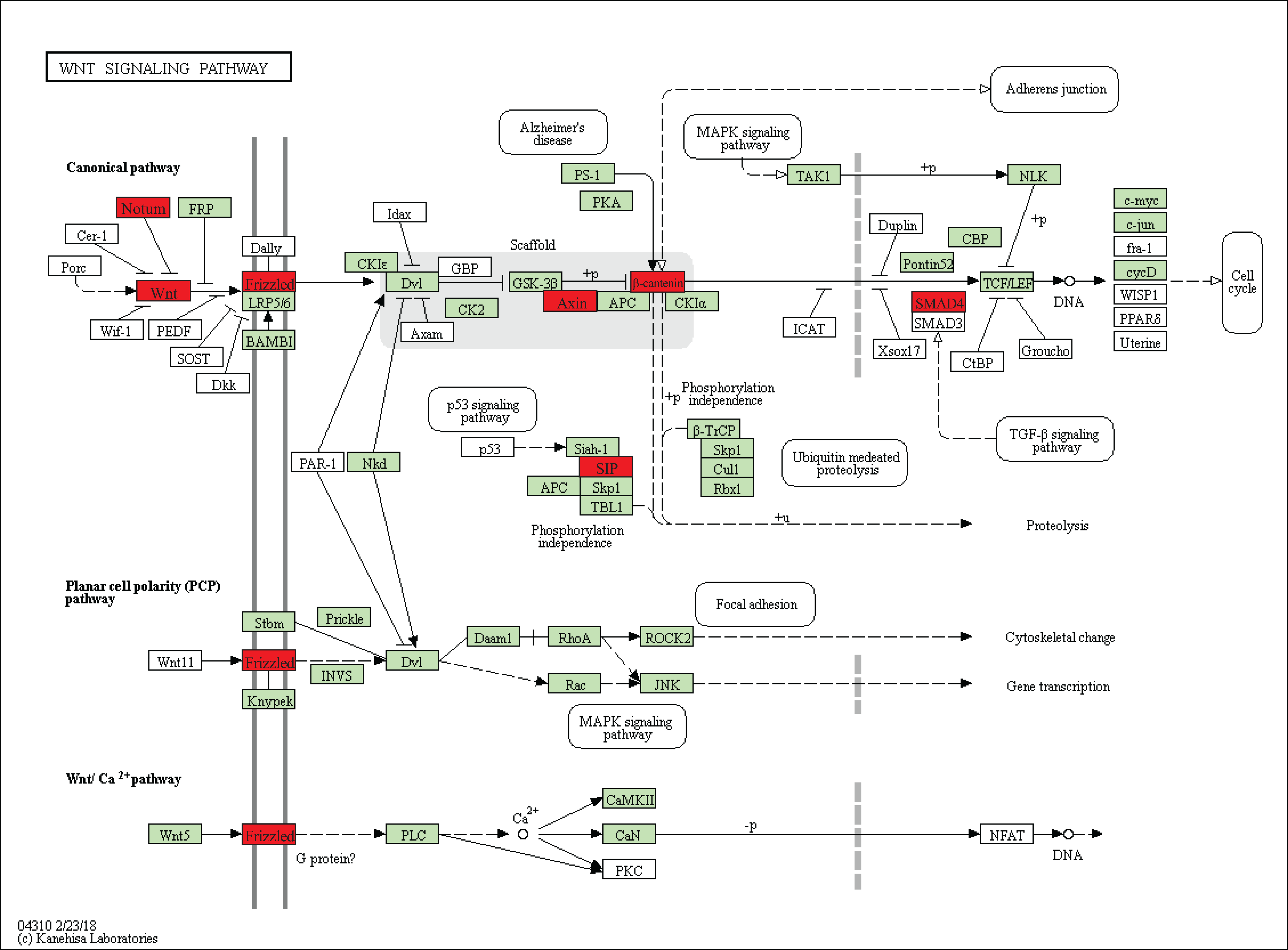
Pictorial representation of the KEGG pathway for WNT signaling. The red shaded boxes indicate the genes having fixed amino acid differences between *A. cervicornis* and *A. palmata*. Green indicates the genes that were found in these genomes but did not differ between the species. White indicates the genes that were not found in the three Acroporid genomes.

The strongest support from the KEGG analysis was for an enrichment of fixed amino acid differences in 12 of 67 ABC transporters (Table 3). The 12 include orthologues of the following three ATP-binding subfamily members: member 7 of subfamily B, member 2 of subfamily D (ABCD2), and member 2 of subfamily G. Judged by the level of inter-species sequence conservation around the variant position, ABCD2 stands out. ABCD2 transports fatty acids and/or long chained fatty acyl-CoAs into the peroxisome (Andreoletti et al. 2017). The variant valine (V) appears to at the beginning of transmembrane helices 3 that is conserved in the majority of coral species, including *A. digitifera* and *A. cervicornis* (Figure S8). In *A. palmata* and *A. millepora* the V is replaced by isoleucine (I). However, the residues predicted to stabilize ABCD proteins and facilitate transport across the membrane are conserved between all corals and the human orthologue (Andreoletti et al. 2017). In vertebrates, the “motif’ SVAHLYSNLTKPILDV is essentially conserved in all mammal, bird, and fish genomes available at the UCSC browser (Figure S9). The only three substitutions pictured in Figure S9 are a somewhat distant I⟶V in hedgehog and rabbit, and V⟶I in opossum at the position variant in *A. palmata*. This extreme level of inter-species protein conservation suggests that the ABCD2 orthologue may function somewhat differently in *A. palmata* and *A. millepora* compared to most other corals. However, the ease with which V and I can be interchanged in nature, because of their biochemical similarity and illustrated by the mammalian substitutions mentioned above, tempers our confidence in this prediction. Still, the apparent near-complete conservation of this particular valine in evolutionary history lends some weight to the hypothesis.

### Genomic Stretches of SNVs

Rather than restricting the analyses to only the fixed SNVs, a larger set of the high-quality SNVs related to the species differences can be identified by interrogating the joint allele-frequency spectrum of the two species. An advantage of this approach over considering justamino acid variants is that it can potentially detect functional changes in non-coding regions, such as promoters or enhancers. We identified 12,279 intervals of consecutive SNVs with high *F*_ST_ values. The genomic intervals ranged from 5 b (NW_015441140.1:321,729-321,734, 4 SNVs with average *F*_ST_ = 1.0) to 27 kb (NW_015441096.1: 814,882-842,464, 8 SNVs with average *F*_ST_ = 0.9217). The top scoring interval covers a 14 kb window in positions 64,603-78,897 of scaffold NW_015441181.1 (Table S5 and Figure 5A). The average *F*_ST_ for the 241 SNVs in this interval is 0.9821, while the average *F*_ST_ for all of the roughly 1.7 million SNVs is 0.1089. Within this interval, there are three gene models: methyltransferase-like protein 12 (MTL12; NCBI: LOC107339088), Wnt inhibitory factor 1-like protein (WIF1; NCBI: LOC107339060), mucin-5AC-like protein (MUC5AC; NCBI: LOC107339062) (Figure 5A).

**Figure 5.**
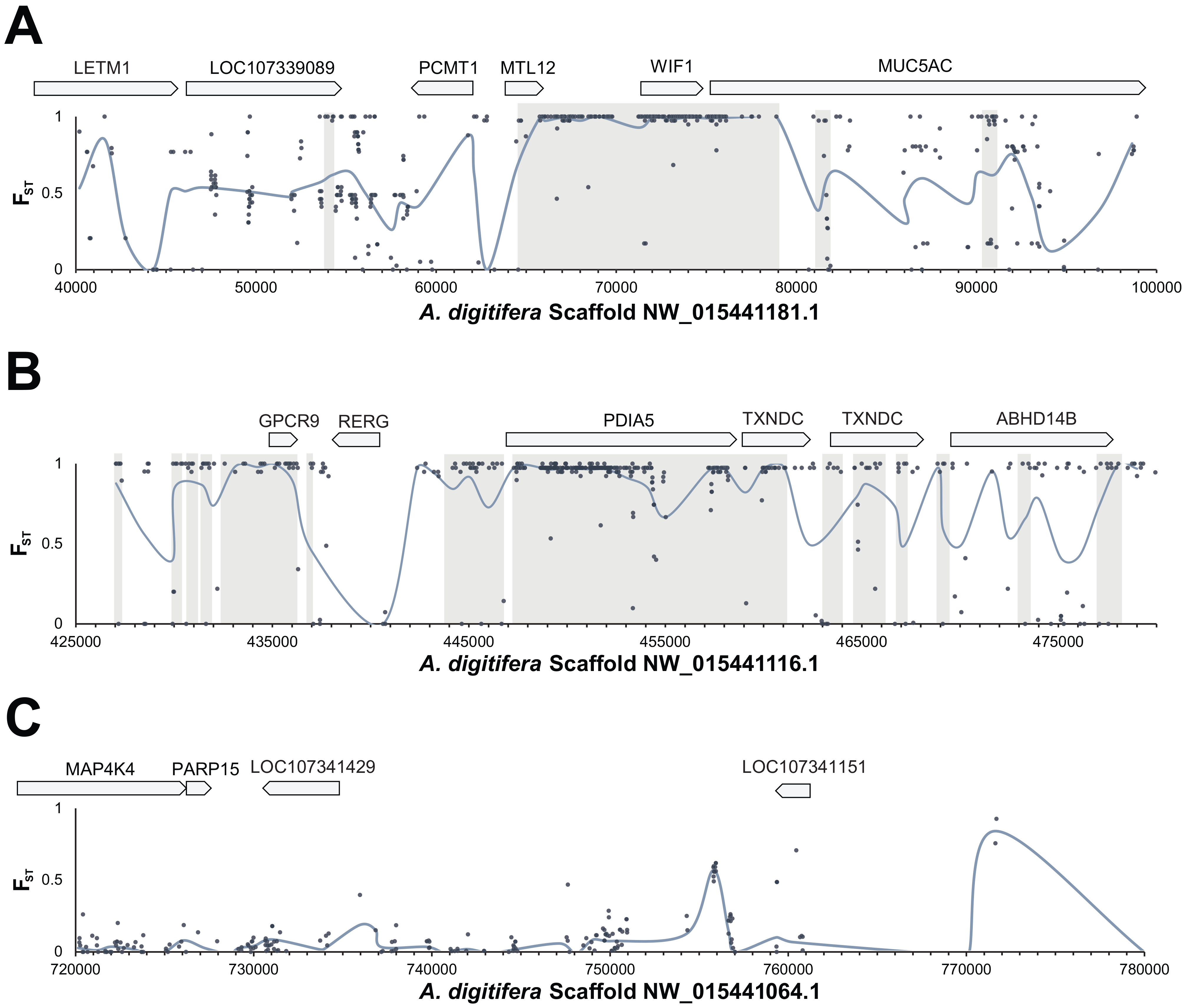
Genomic intervals with or without regions of differentiation between *A. palmata* and *A. cervicornis*. Inter-species allelic differentiation (*F*_ST_) was calculated using the unbiased Reich-Patterson estimator (Reich et al. 2009). Intervals of high scoring SNVs were identified by subtracting 0.90 from each SNV *F*_ST_ value and totaling the score of consecutive SNVs until the score could no longer be increased by an additional SNV on either end. High scoring regions are shaded in light grey along 60 kb genomic windows for the top two scoring intervals, scaffold NW_015441181.1 (A) and scaffold NW_015441116.1 (B), compared to 60 kb genomic window on scaffold NW_015441064.1 with no intervals (C). Grey points are the *F*_ST_ estimate for each SNVs and blue line is the average *F*_ST_ calculated over 1 kb sliding window analysis. Predicted genes within these windows are shown above the graph in grey arrows. In order, genes include mitochondrial proton/calcium exchanger protein (LETM1), *A. digitifera* LOC107339089, protein-L-isoaspartate(D-aspartate) O-methyltransferase (PCMT1), mitochondrial methyltransferase-like protein 12 (MTL12), Wnt inhibitory factor 1 (WIF1), mucin-5AC-like (MUC5AC), G protein-coupled receptor 9 (GPCR9), Ras-related and estrogen-regulated growth inhibitor (RERG), protein disulfide-isomerase A5 (PDIA5), thioredoxin domain containing protein (TXNDC), protein ABHD14B (ABHD14B), mitogen-activated protein kinase kinase kinase kinase 4 (MAP4K4), poly(ADP-ribose) polymerase family member 15 (PARP15), *A. digitifera* L0C107341429, and *A. digitifera* L0C107341151.

The next highest scoring run of high *F*_ST_ values is the 15 kb interval in positions 447289-462570 of scaffold NW_015441116.1 (Figure 5B). The 306 SNVs in this region have an average *F*_ST_ = 0.9756. The most recent NCBI gene annotations mention two intersecting genes in the interval, protein disulfide-isomerase A5-like (PDIA5; NCBI: LOC107334364), mapping to the interval 447,296-458,717, and thioredoxin domain-containing protein 12-like (TXNDC12; NCBI: LOC107334366), mapping to 459,123-462,401 (Table S5 and Figure 5B). Adjacent to this interval are three lower scoring intervals also containing a gene annotated as TXNDC12 (NCBI: LOC107334421), mapping to 463,276-467,160 (Table S5 and Figure 5B). The mapping of LOC107334366 shows a strong match to seven exons, but the mapping of LOC107334364 include weakly aligning exons and missing splice signals. LOC107334364 consists of three weakly conserved tandem repeats, and has partial blastn alignments to position 33-172 of human thioredoxin domain-containing protein 12 precursor (GenBank: NP_056997.1). The shorter sequence LOC107334366 has a blastp alignment (E-value 9e-22) to the same region. In the older Reef Genomics dataset for *A. digitifera*, the corresponding gene for L0C107334364 is Acropora_digitifera_14046l. Thus, based on the newer NCBI annotation, there appears to be either a gene or a pseudo-gene in this highly divergent genomic region of *A. digitifera*.

### SNV Markers for Species Identification and Hybrid Assignment

To aid the design of genotyping studies we identified 894 “PCR-ready” SNVs as those that do not have another SNV, indel, or any (interspersed or tandemly duplicated) repeats within 50 bp (Table 2). We call these the “PCR-ready” SNVs, since in theory they are good candidates for amplification in any of the three *Acropora* species. We validated a subset of eight of these PCR-ready SNVs in additional *A. palmata* (*n*=10) and *A. cervicornis* (*n*=9) samples from across the geographic range (Table S2) using a RFLP assay. The eight markers were designed to digest the PCR product at a single nucleotide base present in only one of the two species (Table S3). For example, at locus NW_015441435.1 position 299429, the variable base between the species (GG in *A. cervicornis* and AA in *A. palmata*) provides a unique recognition site in *A. cervicornis* for the restriction enzyme *Hpy*CH4IV (A^CG_T) that results in digestion of *A. cervicornis* PCR product but not *A. palmata* (Figure S11A). We found that our stringent selection of PCR-ready SNVs are in fact fixed in the additional samples surveyed.

We also screened colonies that were morphologically classified as hybrids between *A. palmata* and *A. cervicornis*. We attempted to refine the hybrid classification of colonies into first or later generation hybrid groups based on the proportion of ancestry from each parental species using five microsatellite markers or the above described eight SNV loci. .

Using the SNV makers, the reference F1 hybrids and seven later generation hybrids were heterozygous at all variable sites, whereas the remaining later generation hybrids (*n*=17) genotypes at each site varied depending on the locus (two examples in Figure S11). Similar to the F1 hybrids, the test set of hybrids were also heterozygous at all loci. For each locus, genotypes were scored to produce a multi-locus genotype (MLG) for each individual.

The congruency of taxon classification was compared between the SNV MLGs and microsatellite MLGs using a discriminant factorial correspondence analysis (DFCA) for each marker set (Figure 6). All *A. cervicornis* samples but one were correctly identified to their taxonomic group using the microsatellite MLGs, but in only 50% of *A. palmata* colonies did the microsatellite clustering coincide with the previous taxon assignment (Table S2 and Figure 6A). In contrast, because of stringency in selecting the fixed SNV loci, there was 100% agreement of the previous taxon assignment of the parental species colony and its SNV MLG classification (thus data points for pure bred samples are overlaid by the group centroid in Figure 6B).

**Figure 6.**
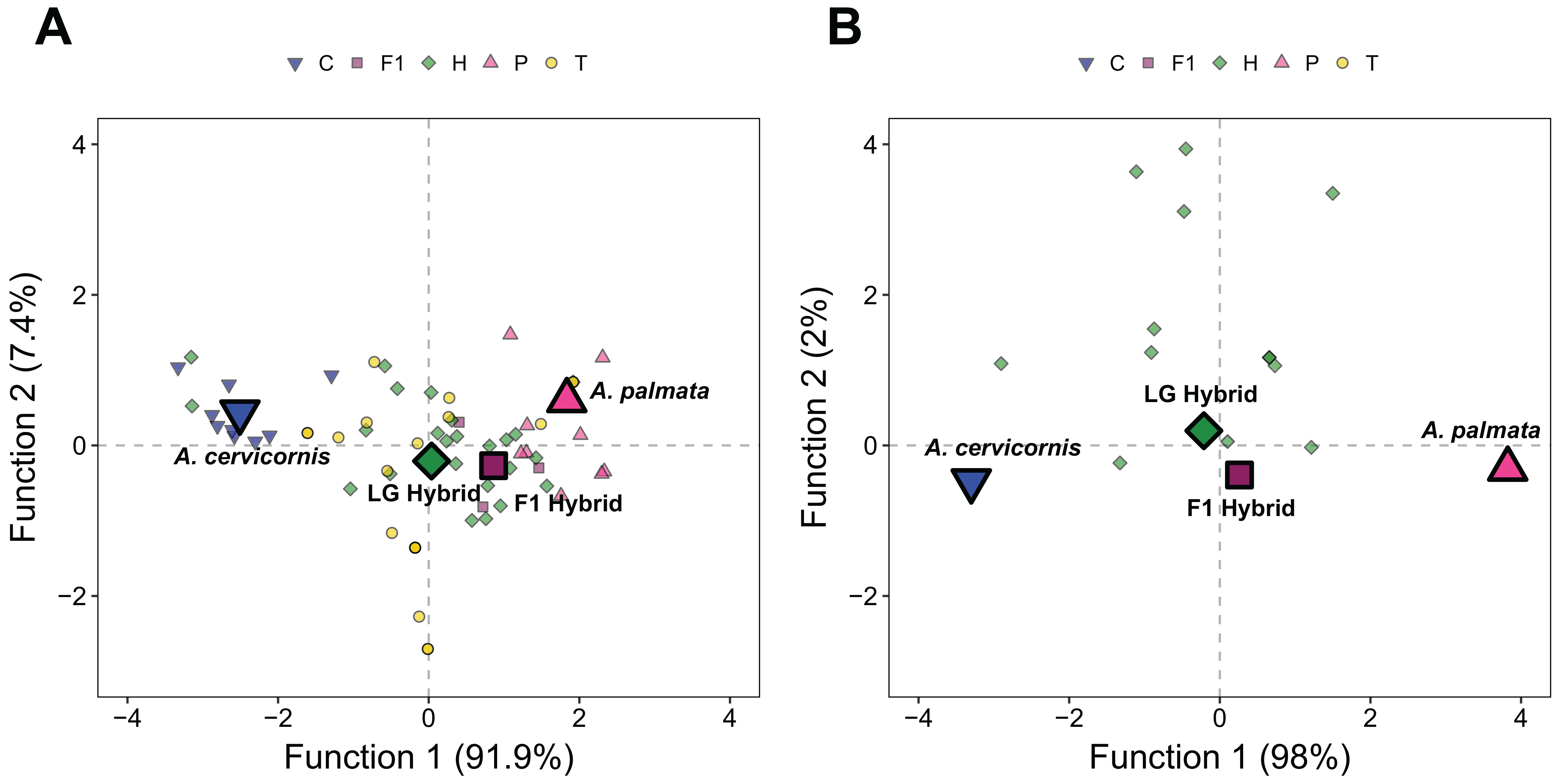
Discriminant factorial correspondence analysis of five microsatellite markers (A) and eight species-specific SNV loci (B). Samples were assigned to four different groups based on their previous taxon assignment: 1. *A. cervicornis* (*n*= 9, blue upside down triangles), 2. *A. palmata* (*n*=10, pink triangles), 3. F1 hybrids (*n*=3, purple squares), and 4. later generation hybrids (*n*=24, green diamonds). The remaining hybrid samples (*n*= 20, yellow circles) had no previous hybrid assignment and acted as our test set for the analysis. The large shapes for each group represent the group centroid, or mean. In panel B, data points for pure bred colonies are not visible because their coordinates are identical to their respective group centroids. F1 hybrids, test hybrids and seven later generation hybrids are also masked as they share the same coordinates as the F1 centroid, representing F1-like hybrids in the data set.

No hybrid samples (F1, later generation or those in the test set) were assigned with high probability to the F1 group with either maker set in the DFCA (Table S2). However, we found that the SNV MLGs of F1 hybrids, seven later generation hybrids and all test hybrids shared the same discriminant function coordinates as the F1 centroid, representing F1-like hybrids in the data set (overlaid by F1 group centroid in Figure 6B). The remaining later generation hybrids were classified as either *A. cervicornis* (*n*=6), *A. palmata* (*n*=1) or hybrid (*n*= 10; Figure 6B and Table S2).

## DISCUSSION

In this study, we have identified inter- and intra-species SNVs and indels between three *Acropora* species. These variants can cause amino acid substitutions that might ultimately alter protein function between these corals. We provided examples of genes with putative fixed-differences between the Caribbean Acroporid species, grouped variants by their KEGG pathways, highlighting examples from the Wnt and ABC transporter pathways, identified highly diverged genomic regions between them and developed a RFLP assay to distinguish species and hybrids. Genomic resources and workflows are available on Galaxy allowing researchers to reproduce the analyses in this paper and apply them to any Acroporid species or other non-model organisms.

### Candidate Loci in Growth and Development

Genes in the Wnt pathway are critical for pattern formation, tissue differentiation in developing embryos and tissue regeneration of Cnidaria (Guder et al. 2006). Interestingly, we found that genes in the Wnt pathway are enriched in fixed amino acid substitutions and an antagonist of this pathway, WIF, has consecutive SNVs with high *F*_ST_ values between *A. cervicornis* and *A. palmata*.

Wnt genes function in primary body axis determination in *Hydra* and *Nematostella* (Hobmayer et al. 2000; Kusserow et al. 2005), and in bud and tentacle formation in *Hydra* (Philipp et al. 2009). Changes in the expression of Wnt genes under high temperatures are hypothesized to result in disassociating *A. palmata* embryos and planulae with bifurcated oral pores, indicating the critical role of this pathway in the ability of coral larvae to develop properly under thermal stress (Polato et al. 2013). The genomic differences in the Wnt genes between elkhorn and staghorn corals reported here (Figure 4) could reflect developmental or growth adaptations that may be influenced by temperature, underscoring the warning that changing ocean temperature can alter the development of corals.

The Wnt pathway continues to regulate coral growth beyond early developmental life stages. In the two Caribbean Acroporids, expression of Wnt genes was higher in the tips of colonies than the base of colonies (Hemond et al. 2014). Differential expression of WIF was not observed in the comparison of the distinct branch regions within or between species or under larval thermal stress in *A. palmata* (Polato et al. 2013; Hemond et al. 2014), but WIF expression in *A. digitifera* did change across the transitional life stages of blastula, gastrula, post-gastrula and planula (Cruciat and Niehrs 2013; Reyes-Bermudez et al. 2016).

Another candidate gene STRADα (Figure 3) is part of the AMP-activated protein kinase (AMPK) pathway, which plays a key role in cellular growth, polarity and metabolism. Under starvation or stressful conditions, the AMPK pathway senses cell energy and triggers a response to inhibit cell proliferation and autophagy (Hawley et al. 2003). Recently, the switch towards activation of AMPK-induced autophagy over apoptosis has been proposed to enhance disease tolerance in immune stimulated corals (Fuess et al. 2017). In this study, STRADα was found to have two non-synonymous mutations and an indel between *A. cervicornis* and *A. palmata* (Figure 3). Although these changes do not occur in a reported site of activity, we cannot ignore the possibility that they are relevant in the interaction of STRADα with MO25Aα and LKB1. The products of these three genes interact together to regulate the AMPK cascade, with STRADα being key for LKB1 protein stability. The extent to which AMPK more broadly contributes to the development and disease tolerance of elkhorn and staghorn corals needs to be further explored.

### Candidate Loci for Microbe Interactions and Cellular Stress

We highlighted several genes with fixed differences between the two Caribbean Acroporids that are involved in innate immunity, membrane transport and oxidative stress in cnidarians. These genes are also important for mediating interactions between the coral host and their microbial symbionts. Corals mediate interactions with foreign microbes by either creating physical barriers or initiating an innate immune response (Palmer and Traylor-Knowles 2012; Oren et al. 2013). Innate immunity is not only activated for the removal of threatening microbes, but also facilitates colonization of beneficial microorganisms within the coral host.

As one of the physical barriers, corals secrete a viscous mucus on the surface of their epithelium that can trap beneficial and pathogenic microbes (Sorokin 1973; Rohwer et al. 2002). Microbial fauna of the mucus can form another line of defense for their host, with evidence that mucus from healthy *A. palmata* inhibits growth of other invading microbes and contributes to the coral antimicrobial activity (Ritchie 2006). This mucus is composed of mucins, one of which might be mucin 5AC that was found to span three divergent genomic intervals between *A. palmata* and *A. cervicornis*. Mucin-like proteins have been found in the skeletal organic matrix of *A. millepora* (Ramos-Silva et al. 2014) and are differentially expressed in the tips of *A. cervicornis* during the day (Hemond and Vollmer 2015) suggesting a potential role for these large glycoproteins in biomineralization as well. Thus, the divergence of mucin in elkhorn and staghorn corals could underlie difference in the composition of their mucus and/or calcification patterns.

Beyond the mucus layer, corals and other cnidarians have a repertoire of innate immune tools to recognize microbial partners from pathogens and remove the latter. The transcription factor NF- κB is one of these tools that regulates expression of immune effector genes, including mucin mentioned above (Sikder et al. 2014). We identified two fixed SNVs in NKIRAS2, an inhibitor of NF-κB transcription (Chen et al. 2004). The two substitutions within this gene were both unique to either *A. palmata* or *A. cervicornis* and neither were shared by the Pacific Acroporids. While the role of NKIRAS1 and −2 are largely unexplored in non-mammal animals, NKIRAS1 has been reported to be one out of nine genes down-regulated at high temperatures in *A. palmata* (Polato et al. 2013).

As a way to interact and exchange nutrients with their beneficial microbes, corals can use ABC transporter proteins. In general, ABC transporters encode for large membrane proteins that can transport different compounds against a concentration gradient using ATP. More specifically, they can transport long-chain fatty acids, enzymes, peptides, lipids, metals, mineral and organic ions, and nitrate. ABC transporters were enriched in fixed amino acid differences between *A. palmata* and *A. cervicornis* (Table 3). Previous characterization of the proteins embedded in a sea anemone symbiosome, the compartment where the symbionts are housed, found one ABC transporter which could facilitate movement of molecules between partners (Peng et al. 2010). ABC transporters were upregulated in response to high CO_2_ concentrations (Kaniewska et al. 2012) and during the day (Bertucci et al. 2015) in *A. millepora* suggesting diverse roles for these proteins, transporting both molecules from the environment and metabolites from their symbionts.

Within the ABC transporters, we analyzed in detail the non-synonymous mutations in ABCD2 between *A. palmata* and *A. cervicornis* (Figure S8). This analysis was limited by the availability of sequences, but allowed us to conclude that the amino acid substitution, though expected to not produce a large functional change, is embedded in a-well conserved motif. The ABCD2 product is involved in the transport of very long-chain acyl-CoA into peroxisomes for β-oxidation. It has been reported that *A. palmata* larvae derive their energy by this mean and that high temperatures induce a change in expression of genes associated with peroxisomal β-oxidation (Polato et al. 2013). This is thought to indicate that larvae of *A. palmata* catabolize their lipid stores more rapidly at elevated temperatures (Polato et al. 2013). Increased lipid catabolism in turn drove the need for additional redox homeostasis proteins to deal with reactive oxygen species (ROS) produced during oxidation of fatty acids (Polato et al. 2013).

Superoxide dismutase, PDIA5 and TXDNC12 are involved in ROS stress-response and antioxidant defense to deal with the oxygen radicals that are produce via the coral host or its symbionts. It has been reported that the antioxidant protein SOD, which converts superoxide anions to hydrogen peroxide, is important to reduce the ROS produced by the coral host and also its dinoflagellate symbiont (Levy et al. 2006), particularly under high temperature stress (Downs et al. 2002), high photosynthetically active radiation (Downs et al. 2002) and salinity stress (Gardner et al. 2016). The genes PDIA5 and TXDNC12 also regulate oxidative stress as well as protein folding. They are both localized to the endoplasmic reticulum and belong to the thioredoxin superfamily of proteins (Galligan and Petersen 2012). These genes were found to span the longest interval of significant genomic differentiation between the two Caribbean species (Figure 5). Thioredoxin-like genes have been differentially expressed in a number of thermal stress experiments on Pacific Acroporids (Starcevic et al. 2010; Souter et al. 2011; Rosic et al. 2014) providing strong support for their role in mediating redox stress. Future research is required to validate the functional consequences of the substitutions in the loci that differ between *A. palmata* and *A. cervicornis* and their putative roles in host cellular stress response, microbial interactions and/or nutrient exchange.

### Mitochondrial SNVs

Unlike other metazoan mitochondrial DNA (mtDNA), cnidarian mtDNA evolves much slower and is almost invariant among conspecifics (van Oppen *et al.* 1999; Shearer *et al.* 2002). However, the so-called control region can be hypervariable compared to the other mtDNA regions in corals (Shearer et al. 2002), and is where the majority of the mitochondrial SNVs in these taxa were identified (Figure S3). The variability in this gene-free region has been used in previous studies to reconstruct the phylogenetic relationship all Acroporid species (van Oppen et al. 2001) and as one of the markers to determine gene-flow between *A. palmata* and *A. cervicornis* from hybridization (Vollmer & Palumbi 2002, 2007). The lack of fixed-differences between the mtDNA of these two species suggests that mito-nuclear conflict might be limited or non-existent during hybridization of these species.

### Species-Specific Diagnostic Markers

We validated eight of the PCR-ready fixed SNVs in additional Acroporid samples and classified the two Acroporid species and their hybrid based on the MLGs of these makers and five microsatellite loci (Figure 6). Currently, microsatellite makers are routinely used to identify Acroporid genotypes and clone mates, but only one of these is a species-specific marker (locus 192) between the Caribbean Acroporid (Baums et al. 2005; Baums et al. 2009). While previous studies have used labor intensive Sanger-sequencing of one mitochondrial and three nuclear loci to study Caribbean hybrid *Acropora* (Van Oppen et al. 2000; Vollmer and Palumbi 2002), PCR-ready fixed SNV markers provide an alternative for high-throughput genotyping and hybrid classification. The detection of only one variable base at each SNV locus can lower genotyping error, avoid difficulties in interpreting heterozygous Sanger sequences and increase reproducibility across labs (Anderson and Garza 2006). Our results indicate a small number of fixed SNVs can outperform the microsatellite makers for taxonomic classification of the species but not necessarily the hybrids. Our inability to discriminate the F1-like hybrids from the later generation hybrids with the DFCA is likely due to the low sample size of reference F1 hybrids (*n*=3). In the case of the SNV markers, the identical MLGs between the F1 hybrids and seven later generation hybrids further reduced our ability to separate the groups. Therefore, with the limited number of PCR-ready SNVs tested, there was no difference in the performance of microsatellite to SNV loci for refining hybrid classification. These results however, indicate that the genomes provide a rich source for PCR-ready SNVs albeit a larger number of SNVs then tested here will need to be assayed before Caribbean Acroporid hybrids can be classified confidently.

## CONCLUSION

By using the genome assembly of *A. digitifera*, we were able to detect differences between *A. cervicornis* and *A. palmata* at various levels, from a single nucleotide substitution to hundreds of nucleotide substitutions over large genomic intervals. We identified genetic differences in key pathways and genes known to be important in the animals’ response to the environmental disturbances and larval development. This project can work as a pilot to gather intra- and interspecies differences between *A. cervicornis* and *A. palmata* across their geographic range. Ultimately, gene knock-down and gene editing experiments are needed to test whether these and other genetic differences have functional consequences and thus could be targets for improving temperature tolerance and growth of corals.

## WEB RESOURCES

The SNV and indel calls for both the nuclear and mitochondrial genomes are available at the Galaxy internet server (usegalaxy.org) (Afgan et al. 2016).

## DATA AVAILABILITY

NCBI Accession numbers for the raw reads are SRR7235977-SRR7236038. Supplemental figures and tables are uploaded to figshare (link).

## ACKNOWLEDGMENTS

This study was funded by NSF OCE-1537959 to IBB, NF, and WM. Thanks to the PSU genomics facility for performing the sequencing. Additional thanks to Meghann Devlin-Durante for assistance with DNA extractions and Macklin Elder for help with the RFLP assay. Samples were collected and exported with appropriate permits.

## AUTHOR CONTRIBUTIONS

AR provided SNV and indel calls and produced the figure of mitochondrial variants. SK extracted the coral DNA, contributed to the analysis of the SNVs and developed and analyzed the RFLP assay. OB generated 3D protein model and performed KEGG pathway enrichment analysis. RB made the variants available on Galaxy. NF provided samples for the genome sequencing and RFLP validation. AR, SK, OB, WM and IBB wrote the paper. The project is being managed by IBB.

## SUPPLEMENTAL DATA

**Table S1. Alignment summary statistics for the various samples included in this study. Each row corresponds to a sample.** The column ‘Generated Reads’ refers to the number of sequences generated for the sample. ‘Mapped Reads’ refers to the sequences that aligned with a mapping quality > 0, and ‘Properly Paired’ refers to the number of reads that align within the expected distance from their mate. ‘Duplicate Reads’ refers to the number of reads that were flagged as putative PCR duplicates. ‘Aligned Reads’ refers to the number of sequences that were aligned to the *A. digitifera* reference using BWA, We present these statistics at both the sequence and the base level. Samples 1012 and 14120 are the deeply sequenced samples.

**Table S2. Results from the discriminant factorial correspondence analysis for the microsatellite and fixed SNV markers. .** Bold clonal IDs indicate repetitive genotypes and grey rows highlight samples where the probability of membership from the discriminant factorial correspondence analysis differs between the two marker sets. For example, sample 2984 was identified as *A. cervicornis* based on morphology and previous posterior probabilities from a NEWHYBRIDS analysis. The discriminant analysis assigned the sample 2984 with 71.53% probability to the hybrid group using the Msat MLG. This is in contrast to the SNP MLG which classified the sample with 96.7 *%* as being *A. cervicornis* in agreement with both the visual identification and NEWHYBRIDS results.

**Table S3. Summary of the eight fixed SNV markers used to assign hybrids and species.** For each SNV, the left nucleotide matches *A. cervicornis* and the right nucleotide matches *A. palmata*.

**Table S4. Fixed SNV marker gene annotation.**

**Table S5. Gene models identified in the two highest scoring *FST* intervals between the 20 samples of *A. cervicornis* and *A. palmata*.**

**Figure S1. Genome coverage distributions of the 21 *Acropora cervicornis* samples.**

**Figure S2. Distance-based phylogenetic tree of the 42 newly sequenced *Acropora* samples.**

**Figure S3. Locations of 172 SNVs and one indel identified in the mitochondrial genome.**

**Figure S4. Superoxide dismutase alignment highlighting the SNV between *A. cervicornis* and *A. digitifera* (Reef Genomics:12779).**

**Figure S5. Alignment of NF-kappa-B inhibitor-interacting Ras-like protein 1 from Acropora_digitifera_6635 to sequences from other corals and the human orthologue (GenBank: NP_065078.1).**

**Figure S6. STE20-related kinase adapter protein alpha isoform 4 (STRADα) truncated alignment of *A. digitifera* protein Reef Genomics: Acropora_digitifera_13579) with coral sequences and human NP_001003788.1 to highlight fixed SNVs and indel in *A. palmata*.**

**Figure S7. Image of the coverage by sequenced reads around the 12-bp deletion of STRADα, showing unanimous agreement of the species difference, for *A. palmata* (A) and *A. cervicornis* (B).**

**Figure S8. ATP-binding cassette sub-family D member 2 alignment of several coral sequences and human orthologue (GenBank: NP_005155.1).**

**Figure S9. Extreme conservation in vertebrates of the motif SVAHLYSNLTKPILDV in ATP-binding cassette sub-family D member 2 (the human gene is transcribed right-to-left).**

**Figure S10. RFLP results for parental species and hybrids for two fixed SNV loci.** The restriction fragment length polymorphism results of two loci, locus NW_015441435.1: 299429 that cuts *A. cervicornis* (A) and locus NW_015441068.1: 984261 that cuts *A. palmata* (B), are displayed in order from left to right for *A. cervicornis* genome sample 13696, *A. palmata* genome sample 13815, F1 hybrid sample 8939, and three later generation (LG) hybrid samples 4062, 6791, and 1302. Each lane is labeled as either marker= M, uncut PCR product = U, or cut PCR product = C. The LG hybrid 1302 presents both heterozygous (A) and homozygous (B) alleles, whereas LG hybrid 6791 is heterozygous and 4062 is homozygous for both loci.

